# A short noncoding RNA modulates gene expression and affects stress response and parasite differentiation in *Leishmania braziliensis*

**DOI:** 10.1101/2024.05.25.595908

**Authors:** José C. Quilles, Caroline R. Espada, Lissur A. Orsine, Tânia A. Defina, Letícia Almeida, Fabíola Holetz, Angela K. Cruz

**Affiliations:** Department of Cell and Molecular Biology, Ribeirão Preto Medical School, FMRP/USP – University of São Paulo, SP, Brazil; Laboratory of Gene Expression Regulation, Carlos Chagas Institute, Oswaldo Cruz Foundation, Curitiba, PR, Brazil

**Keywords:** Noncoding RNA, *Leishmania*, gene expression, metacyclogenesis, nutritional stress

## Abstract

The protozoan parasite *Leishmania* spp. is a causative agent of leishmaniasis, a disease that affects millions of people in more than 80 countries worldwide. Apart from its medical relevance, this organism has a genetic organization that is unique among eukaryotes. Studies of the mechanisms regulating gene expression in *Leishmania* led us to investigate noncoding RNAs (ncRNAs) as regulatory elements. We previously identified differentially expressed (DE) ncRNAs in *Leishmania braziliensis* with potential roles in the parasite biology and development. Herein, we present a functional analysis of one such DE ncRNA, the 147-nucleotide-long transcript ncRNA97, which is preferentially expressed in amastigotes, the replicative form within mammalian phagocytes. By RT-qPCR the ncRNA97 was detected in greater quantities in the nucleus under physiological conditions and in the cytoplasm under nutritional stress. Interestingly, the transcript is protected at the 5’ end but is not processed by the canonical trypanosomatid *trans*-splicing mechanism, according to the RNA circularization assay. ncRNA97 knockout ^(KO)^ and addback ^(AB)^ transfectants were generated and subjected to phenotypic analysis, which revealed that ncRNA97 impairs the starvation response and differentiation to the infective form. Comparative transcriptomics of ncRNA97^KO^ and parental cells revealed that transcripts encoding amastigote-specific proteins were affected. This pioneering work demonstrates that ncRNAs contribute to the developmental regulatory mechanisms of *Leishmania*.

## Introduction

Leishmaniasis is a human disease that can be caused by more than 20 species of *Leishmania* parasites worldwide. In South America, *Leishmania (Viannia) braziliensis* is the most important etiological agent of mucocutaneous leishmaniasis (QUEIROZ et al., 2012), a severe and morbid tegumentary disease (ROGERS et al., 2011). *Leishmania* parasites are transmitted to humans and other mammals through the bite of an infected female sandfly; thus, the parasite lifecycle demands agile adaptation to hostile environments within both vertebrates and invertebrates. Nutritional stress in the sandfly midgut triggers the procyclic form to differentiate into infective the metacyclic parasite form (metacyclogenesis), which is released into the mammalian dermis during a blood meal (BATES, 2007; SILVA-ALMEIDA et al., 2012). Inside macrophages, the metacyclic form differentiates into the replicative intracellular amastigote, which multiplies and spreads to other cells, causing tissue damage and inflammation (SAXENA et al., 2007).

Compared to other eukaryotes, *Leishmania* kinetoplastids possess a unique genetic organization in which functionally unrelated genes are organized as polycistronic units in 34-36 chromosomes in the absence of canonical promoters. Virtually all mRNAs are polycistronically transcribed by RNA-Pol II and cotranscriptionally processed into mature mRNAs. This is accomplished by *trans*-splicing of the 39 nucleotide long mini-exon (ME) sequence of the capped spliced leader RNA to the 5’ end of the new transcript, coupled with the polyadenylation of the upstream gene (LeBowitz et al., 1993; Matthews et al., 1994; Ullu et al., 1993). The control of gene expression in these parasites involves coordinated mechanisms that respond to specific environmental triggers during the life cycle (DE PABLOS; FERREIRA; WALRAD, 2016; DE PABLOS et al., 2019). Due to this atypical genome organization, co- and posttranscriptional mechanisms are critical for the control of gene expression in these protozoans (CLAYTON, 2014, 2016; MARTÍNEZ-CALVILLO et al., 2018). The discovery of noncoding RNAs (ncRNAs) that control gene expression in different organisms has raised the question of whether ncRNAs could play a similar role in trypanosomatids. Aside from the housekeeping ncRNAs (tRNAs, rRNA, and snRNAs) (ERNST; MORTON, 2013), ncRNAs have diverse functions and can be processed in a range of different ways (CHAN; TAY, 2018; SHIH et al., 2020). An arbitrary classification of long (> 200 nt) and short (< 200 nt) sequences is frequently used, but specific characteristics, such as genomic localization, molecular interactions, and functions, can also be used (ERNST; MORTON, 2013). However, a recent novel classification defines long ncRNAs as those >500 nucleotides long (MATTICK et al., 2023). The best studied short noncoding RNAs are microRNAs (miRNAs), which act as *cis-* or *trans-*elements by binding to the 3’-untranslated region (UTR) of a target mRNA and modulating its stability and/or translation (GUO et al., 2010). Despite their short length of only 22 nucleotides, miRNAs play a major role in gene regulation, and their misregulation is connected to a range of human diseases (GHILDIYAL; ZAMORE, 2009; FERNANDES et al., 2019).

Although it has been 30 years since the first functional eukaryotic ncRNA was discovered (BRANNAN et al., 1990), only a few ncRNAs have been detected in trypanosomatids (IRIBAR; TOSI; CRUZ, 2003; DUMAS et al., 2006; ZHENG et al., 2013; FREITAS CASTRO et al., 2017; CHIKNE et al., 2019; GUEGAN et al., 2022). A small nucleolar ncRNA regulates differentiation in *Trypanosoma brucei* parasites by manipulating the expression of two essential differentiation factors (GUEGAN et al., 2022). In *Leishmania* parasites, the ncRNA *ODD3* emerges from the 3’UTR of only one of the gene copies encoding the ribosomal protein S16 (FREITAS CASTRO et al., 2017), but its function remains unknown. We recently identified approximately 3,600 ncRNAs that are differentially expressed (DE) in different developmental stages of *Leishmania braziliensis* (RUY et al., 2019) in an effort to understand the possible functions and relevance of these ncRNAs in life cycle regulation. Here, we characterized one of these DE ncRNAs and showed that it modulates the expression of a group of genes and that it is involved in the regulation of the nutritional stress response and in metacyclogenesis, an essential differentiation process for parasite development.

## Results

### Confirmation, differential expression and processing feature analysis of ncRNA97

RNA-seq predicted ncRNA97 to be a 147-nucleotide-long transcript (Figure 1A) preferentially expressed in amastigotes (RUY et al., 2019). It is located on chromosome 20, and it has been annotated as an independent transcript between two protein-encoding genes (Figure 1A). Nevertheless, the lack of a clear gap in RNA read density between the upstream transcript and the ncRNA97 sequence suggests that ncRNA97 could also be within the 3’UTR of the upstream transcript (Figure 1A). Therefore, we confirmed the differential expression of ncRNA97 by RT‒ qPCR (Figure 1B) and determined its size and the nature of its 3’ and 5’ ends by a circular RNA assay (HANG et al., 2015a). Briefly, total RNA was treated (or not treated) with tobacco acid pyrophosphatase (TAP) to generate 5’-monophosphate RNAs of transcripts bearing a cap to allow circularization after RNA ligase addition. From this circular RNA, cDNA was generated, and primers specific to the ncRNA97 sequence were used, which were directed outward of the transcript in each extremity. Different PCR products were identified (Supplemental Figure 1D), and three bands of approximately 150, 500 and 600 nucleotides were sensitive to TAP and likely to be capped or triphosphorylated RNA (Figure 1C). The TAP-sensitive amplicon of 500 nucleotides and seven other PCR products, ranging from 177 to 838 nucleotides, were sequenced, and the predicted ncRNA97 sequence was confirmed. The poly(A) tail length varied from 0 ≈ 64 nucleotides (Figure 1D and Supplemental Figure 1E). Notably, no ME was detected in any sequence, even in the TAP-dependent PCR product (band F.3 in Supplemental Figure 1E). This was unexpected, as the 5’ cap of trypanosomatid mRNAs is added via the capped ME sequence in the *trans*-splicing reaction. To further confirm the protection at the 5’ end of this transcript, total RNA was treated or not treated with mRNA decapping enzyme (MDE) plus 5’→3’ XRN-1 exonuclease to degrade 5’-monophosphate RNA. Afterward, the remaining RNA was reverse-transcribed to DNA, and ncRNA97 was amplified by PCR. Amastin (LBRM2903_080014000) and rRNA45 transcripts were used as coding and noncoding controls, respectively. PCR products were obtained for all transcripts, independent of the treatment (MDE or XRN-1) (Figure 1E), suggesting 5’ protection of these RNAs. After MDE and XRN-1 treatments, rRNA45 and amastin transcripts were barely detected, while ncRNA97 was very evident. Together, these results indicate that ncRNA97 is not a 5’-monophosphorylated RNA but is protected at its 5’ end in a way that is distinct from the traditional CAP found in *Leishmania* coding genes. Additionally, secondary structure in six of the seven ncRNA97 isoforms detected by the circularization assay was predicted to be an elongated hairpin with two large and one smaller loop (Supplemental Figure 2), which might be relevant for the function or stability of the ncRNA97.

**Figure 1.**
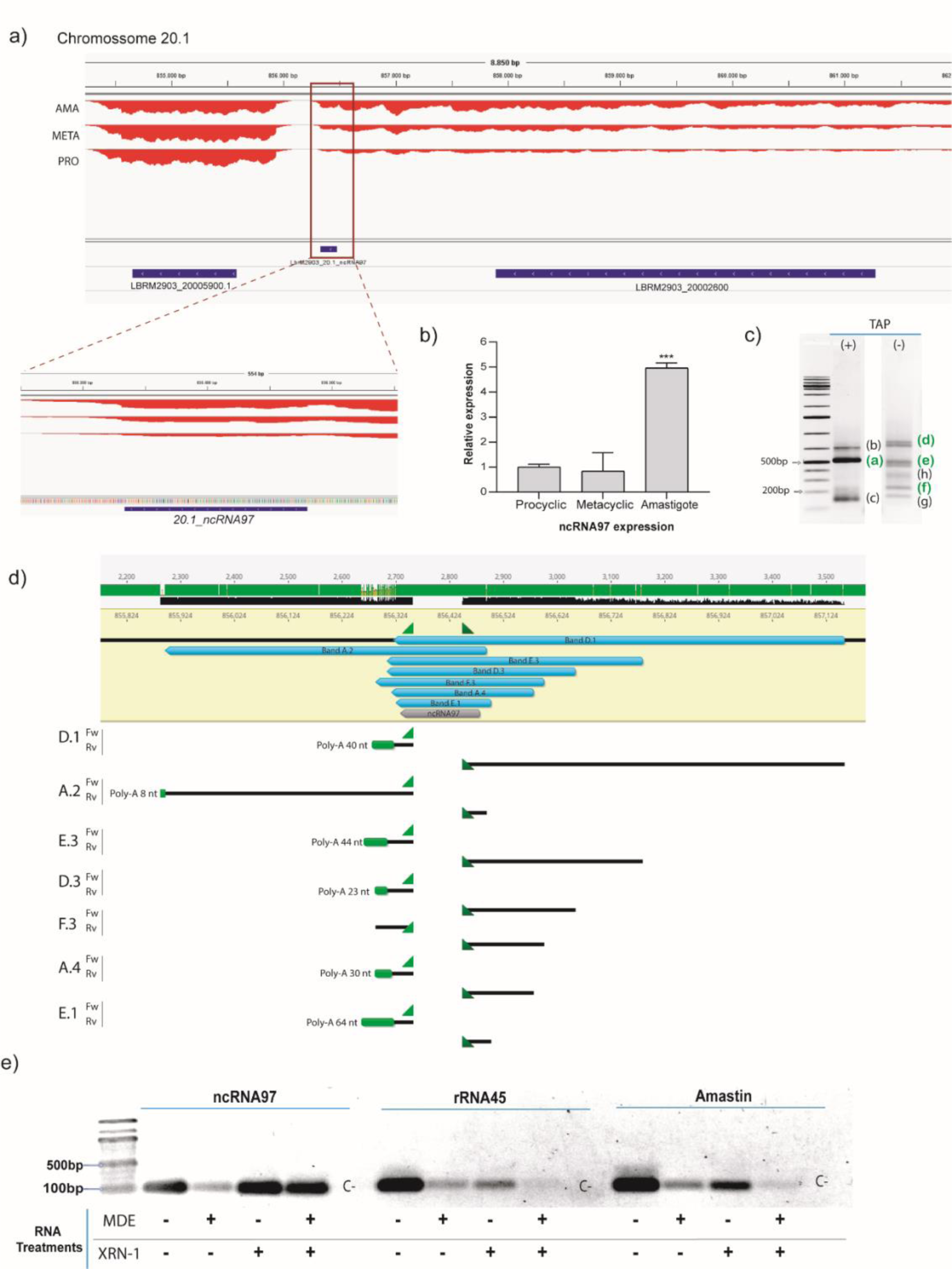
General overview and processing of ncRNA97. (A) Genomic location of ncRNA97 in an intergenic region on chromosome 20.1, with no clear difference in the number of reads in the gap between ncRNA97 and its upstream gene. (B) Confirmation of differential expression of ncRNA97 in axenic amastigotes compared to procyclic and metacyclic amastigotes by RT‒qPCR. (C) RNA circularization assay with or without TAP treatment (see SF1A for methods) and detection of amplified transcript ends by conventional PCR. TAP-dependent and TAP-independent bands were observed, and the bands identified in green were successfully cloned and sequenced. A band of ≈500 bp (a) is a putatively capped transcript, since it is detectable only after decapping treatment. (D) Sequencing revealed different lengths for ncRNA97 transcripts in the cell, with the predicted sequence identified in all transcripts. Poly-A tails are indicated in green. No mini-exon was detected in any of the sequenced ncRNA97 transcripts. (E) Total RNA samples were treated (+) or not (-) with mRNA decapping enzyme (MDE) for cap removal and/or with XRN-1 5’→3’ exonuclease for 5’-monophosphate RNA degradation. Then, the RNA was recovered by chloroform extraction and ethanol precipitation, which was followed by reverse transcription and conventional PCR to detect conventionally 5’ capped transcripts. As already suggested by TAP treatment, ncRNA97 was protected at the 5’ end but not by a conventional trypanosomatids cap. Amastin (LBRM2903_080014000) and rRNA45 transcripts were used as coding and noncoding RNA controls, respectively. A negative control (C-) was generated via PCR with no DNA input.

### ncRNA97 functions: *cis*- and *trans*-regulation of gene expression

ncRNA97 knockout (^KO^) parasites were generated in the *L. braziliensis* M2903 strain via CRISPR/Cas9 genome editing (BENEKE et al., 2017), and homozygote parasites were confirmed by conventional PCR. The genomic position of the primers used for the replacement of ncRNA97 with a selectable marker, the sequence of the guide and donor, and the confirmation of the replacement are shown in Supplemental Figure 3A and 3B. The ncRNA97 knockout negatively affected the transcript levels of the gene upstream (UP) of ncRNA97 (LBRM2903_200026000 – FtsX-like protein), whereas the transcript level of the downstream (DW) gene (LBRM2903_200025900 – hypothetical protein) was not affected (Figure 2A). The localization of the FtsX-like transcript was not altered in ncRNA97^KO^ cells (Figure 2B), indicating that ncRNA97 does not function in regulating nuclear export or cytoplasmic stability. Add-back (ncRNA97^AB^) parasites were generated by transfecting the ncRNA97^KO^ cell line with a plasmid encoding the ncRNA97 sequence (Supplemental Figure 4). RT‒qPCR confirmed that the expression of ncRNA97 in the add-back cells was ∼3 times greater than that in the parental cells (Figure 2C). Importantly, the levels of the downregulated FtsX-like gene fully recovered to the levels in the parental cells (Figure 2C). The rescue of FtsX transcript levels by ectopic ncRNA97 expression indicates a *trans*-regulatory function for ncRNA97, albeit emerging from its (potential) target, suggesting that it may act as a *cis* ncRNA.

**Figure 2.**
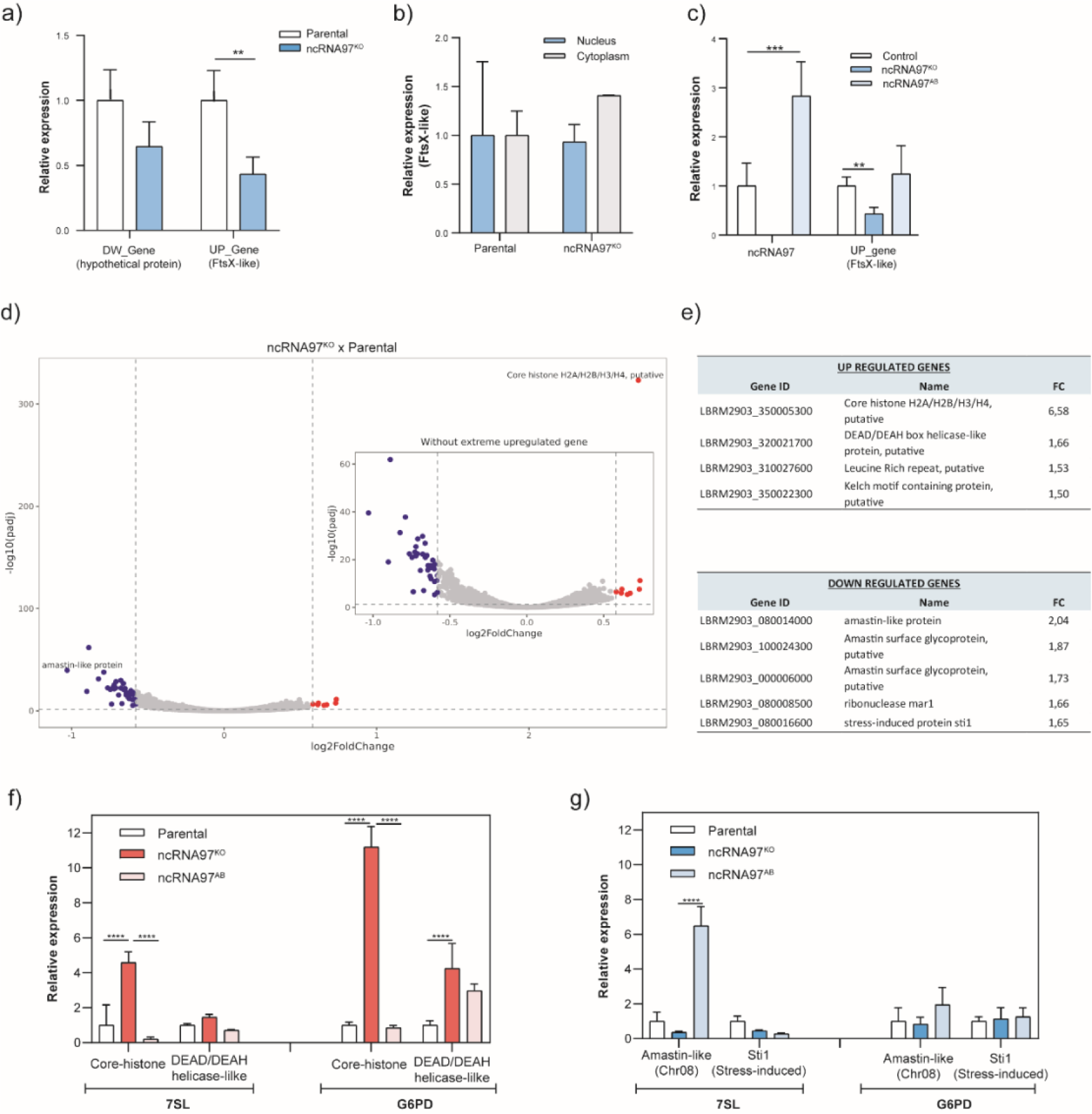
Epigenetic alterations in ncRNA97^KO^ parasites. (A) Downregulation of the upstream gene LBRM2903_20002600 (FtsX-like) in the ncRNA97^KO^ parasites was observed by RT-qPCR. (B) Despite the downregulation, no change in the cellular location of the FtsX-like transcript was observed when the fractionated RNA extracts from parental and ncRNA97^KO^ parasites were compared. (C) The expression level of the upstream gene was rescued to parental levels in ncRNA97^AB^ parasites. (D) Volcano plot showing genes with differential expression between ncRNA97^KO^ and parental cells. (E) The top 4 and 5 genes identified as differentially expressed in the ncRNA97^KO^ cells based on their *p* values for up- and downregulated genes, respectively. (F) The levels of upregulated and (G) downregulated genes levels were restored to near-parental levels in ncRNA97^AB^ parasites. 7SL and G6PD transcripts were used as housekeeping genes to normalize the data.

**Figure 3.**
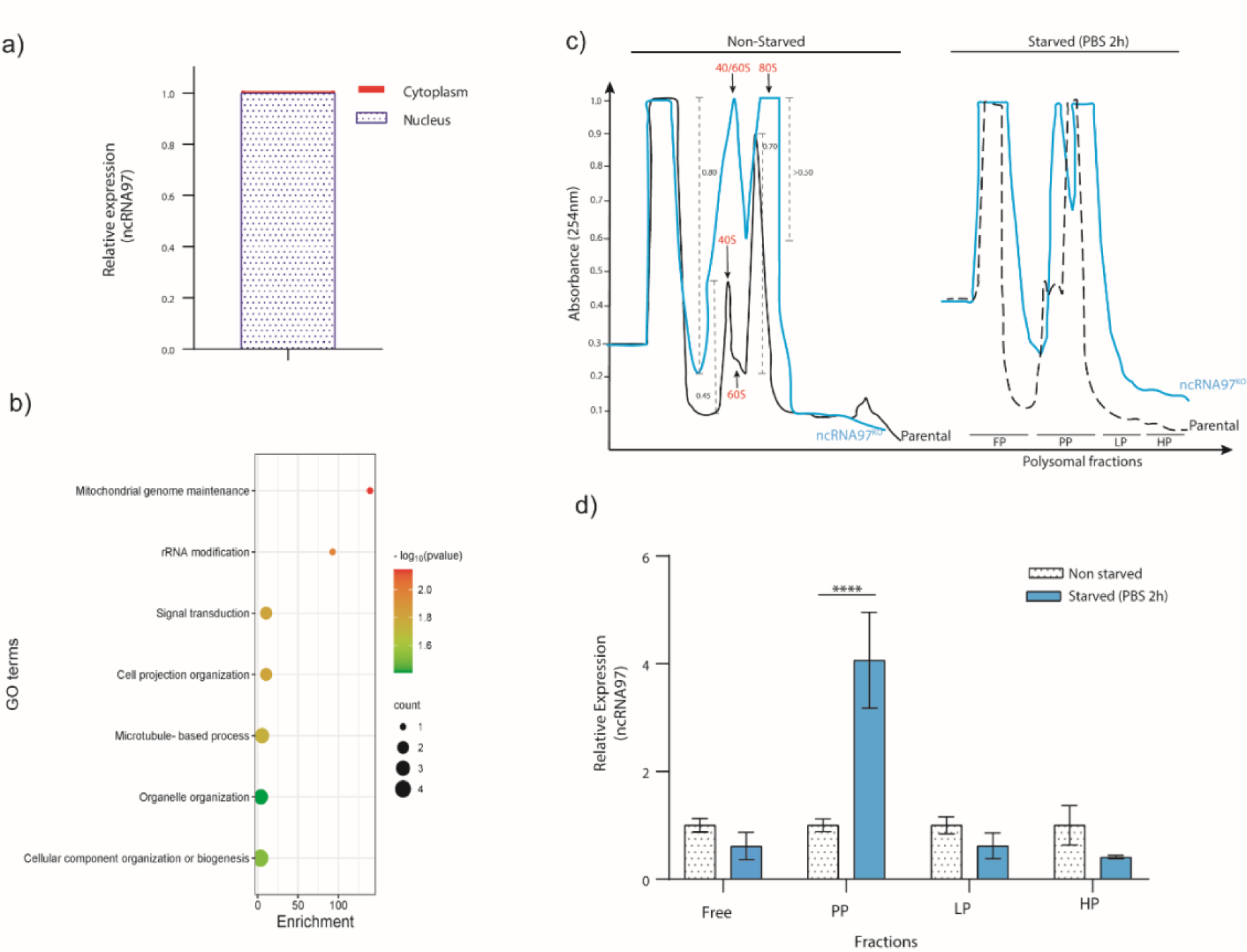
Intracellular localization of ncRNA97 and its respective interacting proteins. (A) Total RNA from promastigotes in the log phase was fractionated, and the relative expression of ncRNA97 was estimated by RT‒qPCR; the standard deviation was lower than 5%. (B) After statistical analysis using a *t* test and considering a *p* value <0.05, the 63 proteins identified as specifically binding to the ncRNA97 sequence by *in vitro* pulldown assay were subjected to GO analysis. See the supplementary material for the complete list. (C) Alterations in the polysome profile of ncRNA97^KO^ parasites under physiological conditions were observed for free ribosomal subunits. (D) Quantification of the ncRNA97 level by RT‒qPCR in the free, pre-polysome (PP), light polysome (LP) and heavy polysome (HP) fractions of parental cells.

Next, to investigate the possible *trans*-regulatory function of ncRNA97, the transcriptome of the parental cells was compared with that of ncRNA97^KO^ parasites (Figure 2D). A relatively small number of genes were upregulated (8) or downregulated (37) (Supplemental Figure 5), among which the top 4/5 (up/down) with the highest fold change (FC) were considered for further investigation. The transcript levels of the putative core histones H2A/H2B/H3/H4 and DEAD/DEAH box helicase were 6.6 and 1.7 times greater, respectively, in the ncRNA97^KO^ cells than in the parental cells (Figure 2E). Interestingly, three out of the five downregulated transcripts encoded amastin, with FCs between 2.0 and 1.7, while the other two encoded the ribonuclease mar1 and stress-induced protein 1 (STI1) (Figure 2E). Amastin, a multigene family with most isoforms exclusively or preferentially expressed in amastigotes, is crucial for the intracellular viability of *L. braziliensis* amastigotes (DE PAIVA et al., 2015). Notably, these three downregulated transcripts encode different amastin isoforms with divergent peptide sequences (Supplemental Figure 6). Additionally, divergence in the untranslated regions of these transcripts might suggest different regulatory mechanisms for their expression (WU et al., 2000). To confirm the RNA-seq results, RT‒qPCR was performed for two up- and downregulated genes. Significant differences between the ncRNA97^KO^ and ncRNA97^AB^ cells were observed for the two upregulated genes, in agreement with the RNA-seq data, but not for the downregulated genes (Figure 2F and 2G, respectively). Importantly, the RT‒qPCR assays were performed in promastigotes, and the low transcript levels of amastin (LBRM2903_080014000) in this stage may have impaired the evaluation of differences, as indicated by the ΔCT values of the RT‒qPCR analysis (Supplemental Figure 3E). However, in ncRNA97^KO^, the ΔCT values were greater than 37, indicating that this gene is expressed at very low levels in ncRNA97^KO^ parasites. Thus, considering their ΔCT values, we may assume that the expression levels of these genes recovered to the parental level in the add-back parasites (Supplemental Figure 3E). These results suggest a correlation between the levels of ncRNA97 and the abundance of certain protein-coding transcripts.

### ncRNA97 protein partners and putative functions

Knowing that ncRNA functions might depend on their intracellular localization and association with proteins (JUNGE et al., 2017), after total RNA fractionation, we quantified the intracellular location of ncRNA97 by RT‒qPCR; we observed predominantly nuclear localization in promastigotes during log phase growth (Figure 3A). Then, we searched for protein binding partners of ncRNA97 by pulldown *in vitro* assays and mass spectrometry. Statistical analysis (*p*<0.05) revealed 63 proteins enriched for interactions with ncRNA97 relative to the aptamer only (Supplementary Material Supplemental Figure 7B). Of these, 25 had predicted functions and were subjected to Gene Ontology (GO) analysis to identify potential processes in which ncRNA97 could be involved (TANG et al., 2023).

Interestingly, one of the identified proteins was a putative rRNA methyltransferase protein (A4HFX5_LEIBR) related to rRNA modification according to its GO annotations (Figure 3B and Supplemental Figure 7A). This protein is homologous to Spb1 from *Trypanosoma brucei,* a protein that is involved in rRNA methylation and processing and relevant to ribosome assembly (Sekulski, 2022). Thus, to investigate any phenotype correlating ribosome and ribosomal protein binding to ncRNA97, we examined the polysome profiles of parental and ncRNA97^KO^ parasites. Despite the poor polysome signal detected for both parental and ncRNA97^KO^ promastigotes, under physiological conditions, the peaks corresponding to the 40S and 60S free ribosome subunits (RSs) were higher in the ncRNA97^KO^ parasites than in the parental cells (Figure 3C). In addition, we compared the ribosomal profiles of the ncRNA97^KO^ and parental lines after nutritional stress induction by PBS. After 2 h in PBS, no substantial alterations in the profile of the ncRNA97^KO^ or parental cell parasites were observed (Figure 3C). Interestingly, RT‒qPCR quantification of ncRNA97 in the fractions collected from different sucrose densities revealed that ncRNA97 accumulated at a density corresponding to the pre-polysome (PP) fraction when the parasites were subjected to PBS starvation (Figure 3D). Remarkably, the ribosomal protein L31, a large ribosomal subunit constituent (LIU et al., 2016) and a eukaryotic translation factor were detected among the ncRNA97 partners in the pulldown assays. The pulldown and polysome profiling results led us to hypothesize that ncRNA97 might play a role in ribosomal processing/assembly.

### ncRNA97 affects growth, metacyclogenesis, and the nutritional stress response

ncRNA97^KO^ parasites were then screened for phenotypic alterations relative to the parental cell line (Supplemental Figure 3F). Compared with control cells, cultured ncRNA97^KO^ cells had a longer doubling time (Supplemental Figure 3G). This growth phenotype was not caused by the downregulation of the FtsX-like transcript, the ncRNA97-upstream gene, as FtsX-like^KO^ cells grew normally (Figure 2C, Supplemental Figure 3G). The increased doubling time of the ncRNA97^KO^ cells may partially explain their poorer metacyclogenesis rate, which was indicated by the results of the metacyclic enrichment Ficoll assay (SPÄTH; BEVERLEY, 2001) (Figure 4A). Additionally, MTT colorimetric assays to evaluate promastigote viability after starvation in PBS for 4 h indicated that ncRNA97^KO^ parasites were more sensitive to nutritional stress than the parental and FtsX-like^KO^ parasites (Figure 4B). Moreover, after 4 h of starvation, the ncRNA97 levels in the parental cells decreased significantly, with partial recovery after 24 h of incubation in fresh medium (Figure 4C). Axenic amastigotes were evaluated for oxidative stress response, and despite the lower resistance of ncRNA97^KO^ axenic amastigotes (Supplemental Figure 3H) to H_2_O_2_, the parasite infectivity in THP-1 macrophages and intracellular proliferation *in vitro* were similar to those in the parental cells (Supplemental Figure 3I and J). The metacyclogenesis rate recovered to the parental level in ncRNA97^AB^ parasites, suggesting that ncRNA97 impairs metacyclogenesis in ncRNA97^KO^ parasites (Figure 4D).

**Figure 4.**
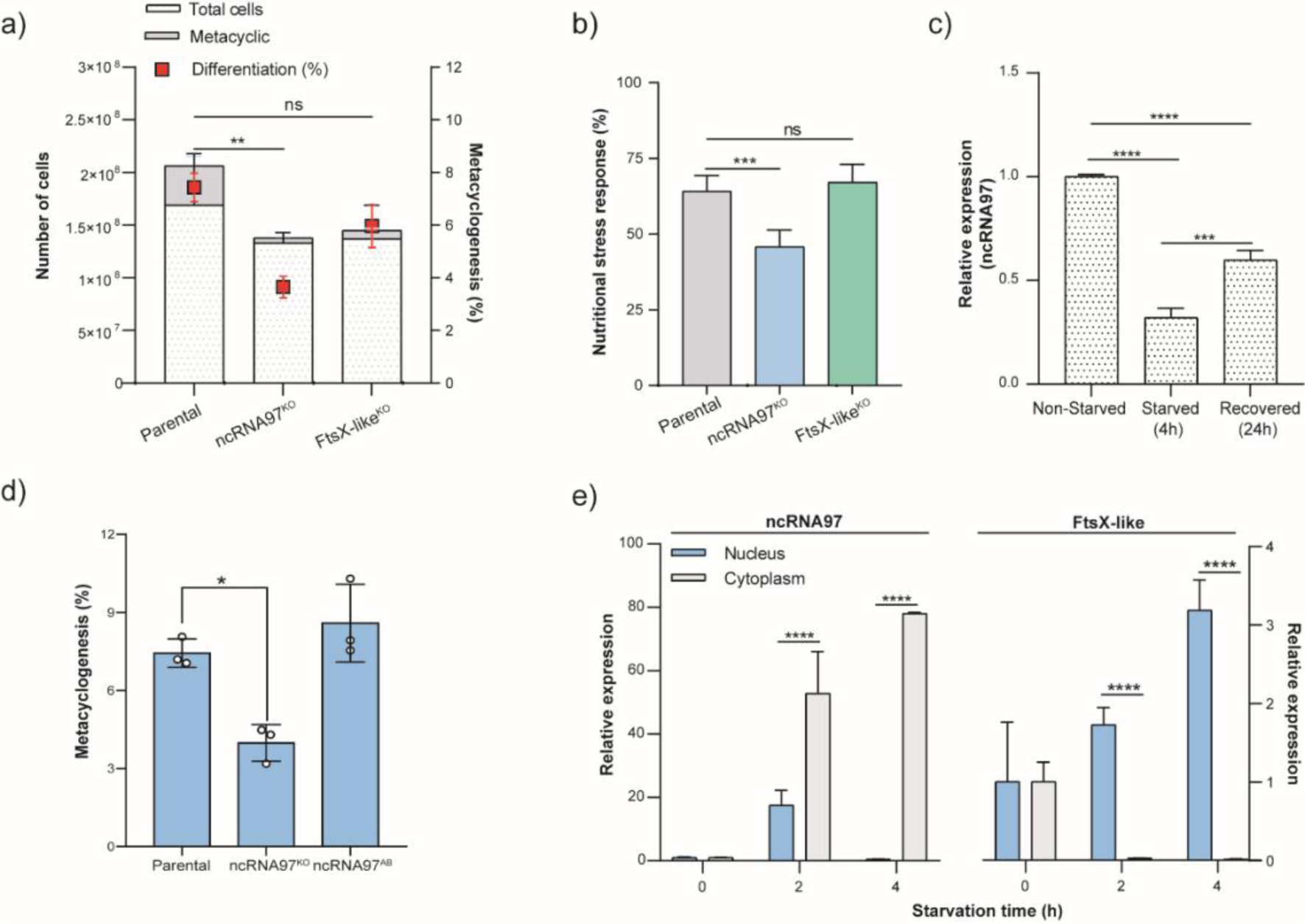
Phenotypic alterations in ncRNA97^KO^ cells and ncRNA97 cell distribution and intracellular localization during starvation. (A) The percentage of the metacyclic form in the stationary phase was used to estimate the metacyclogenesis rate by a Ficoll density gradient assay (Späth and Beverley, 2001), which revealed an impairment in the metacyclogenesis rate exclusively in ncRNA97^KO^ parasites. (B) Parasites lacking ncRNA97 were more sensitive to 4 h of nutritional stress in PBS, based on the MTT viability assay. (C) Transcript level of ncRNA97 determined by RT‒ qPCR showing a decrease during starvation with partial recovery after 24 h. Quantification is relative to the expression levels of the endogenous transcript (G6PDH). (D) The rate of metacyclogenesis in ncRNA97AB parasites recovered to the parental level. (E) Cytoplasmic accumulation of ncRNA97 transcripts increased in a starvation time-dependent manner and was not correlated with the localization of the upstream transcript LBRM2903_20002600, which accumulated in the nucleus.

Next, we evaluated the intracellular distribution of ncRNA97 via subcellular fractionation and subsequent quantification of the nuclear and cytoplasmic levels of ncRNA97 under normal and starvation conditions by RT‒qPCR. Interestingly, both the subcellular distribution and levels of ncRNA97 changed with nutritional stress. The cytoplasmic accumulation of ncRNA97 transcripts increased with starvation time to approximately 80-fold after 4 h of starvation in PBS and was barely detectable in the nucleus under these conditions (Figure 4E). Notably, the observed dynamics of ncRNA97 expression and localization are markedly distinct from those of its upstream gene, since the FtsX-like transcript accumulated in the nucleus after nutritional stress (Figure 4E). These results corroborate the observed increase in ncRNA97 levels in the ribosomal fraction under starvation conditions (Figure 4D). Additionally, no difference in the nuclear or cytoplasmic levels of FtsX-like transcripts was detected between the parental and ncRNA97^KO^ cells (Figure 2B).

## Discussion

Herein, we investigated the functions, partners, and targets of ncRNA97, an ncRNA preferentially expressed in amastigotes, which was selected from approximately three thousand ncRNAs with putative differential expression across developmental stages in *L. braziliensis* (RUY et al., 2019). Our main findings indicate that ncRNA97 is involved in gene expression regulation and affects metacyclogenesis and the parasite response to nutritional stress. We confirmed the presence, differential expression, length, and processing of the ncRNA97 predicted based on RNA-seq data (RUY et al., 2019). RT‒qPCR confirmed its differential expression in the amastigote stage, and the length and polymorphism of the transcripts (varying from 177 to 838 nucleotides) encompassing the computationally predicted sequence were revealed via an RNA circularization assay (Figure 1C). For the seven sequenced ncRNA97 transcripts, we have no data on the preferential isoform or whether they vary depending on the stage or are generated on subsequent processing and cleavages, but we note that all but one transcript possess a conserved secondary structure. We might hypothesize that the observed structure is central to the interaction of ncRNA97 with partner proteins and consequently to its functions, but this hypothesis must be investigated for confirmation.

The RNA circularization assay in association with different enzyme treatments indicated that the different lengths and termini of the transcripts originated from the ncRNA97 genomic region. The approach used permitted the detection of poly(A) tails of variable lengths at the 3’ end in six out of seven cloned transcripts. Additionally, the resistance of ncRNA97 isoforms to 5’→3’ exonuclease treatment indicates that they are not 5’-monophosphate RNAs but might carry an alternative cap structure at the 5’ end.

Based on the biogenesis of 3’UTR-derived ncRNAs from mammalian cells (MERCER et al., 2011), we propose a mechanism in which ncRNA97 is released from the 3’UTR of the upstream mRNA through the action of endoribonucleases (LUCIANO; BELASCO, 2019). To date, we have not conducted any validation assays to confirm that the upstream gene and ncRNA97 are generated as a single transcript and that cleavage to release ncRNA97 occurs at a posterior step to empirically prove this model. Notably, however, the transcriptome profile obtained (Figure 1A) showed no clear difference in the number of reads between the ncRNAs and the upstream mRNAs, supporting this hypothesis. If our model is correct, alternative processing machinery and 5’ protection may occur in *Leishmania.* Several examples of alternative noncanonical RNA capping and different endoribonucleases are available that may be responsible for the biogenesis of ncRNAs (WIEDERMANNOVÁ; JULIUS; YUZENKOVA, 2021; MALKA et al., 2022). Our results demonstrated that ncRNA97, although protected, has no ME sequence at the 5’ end but carries a poly(A) tail at the 3’ end. In future studies, the role of endoribonucleases in the cleavage of these ncRNAs must be investigated.

With no information about the possible roles of the studied DE ncRNA, to test the possible functional effects of ncRNA97, we generated ncRNA97^KO^ parasites, and in this background, we reinserted the transcript sequence to be expressed from a plasmid to generate ncRNA97^AB^ cells. In addition to these strains and the parental line, we also generated a knockout of the ncRNA97 upstream gene FtsX-like, thus ensuring that any phenotypic changes observed were due to ncRNA97 knockout. These cell lines were analyzed for several properties, such as growth under axenic conditions to evaluate phenotypic alterations in promastigote doubling time, *in vitro* infectivity, and metacyclogenesis, as well as for response to nutritional and oxidative stresses, conditions mimicking some of the hostile environments encountered during the parasite lifecycle.

A longer promastigote doubling time (Supplemental Figure 3G) and metacyclogenesis impairment (Figure 4A) compared to those in parental and FtsX-like^KO^ cell lines were observed only for ncRNA97^KO^ parasites, suggesting a cellular effect directly related to ncRNA97. The ncRNA97^KO^ parasite response to starvation was not comparable to that of the parental cell line (Figure 4B), probably due to their slower growth and lower frequency of cell differentiation. However, unlike in procyclic promastigotes, ncRNA97^KO^ intracellular amastigote replication inside macrophages was similar to that in parental cells *in vitro* (Supplemental Figure 3I and J). Confirming the relevance of ncRNA97 for amastigotes, ncRNA97^KO^ parasites were more sensitive than parental parasites to recovery from oxidative stress (24 h in H_2_O_2_) (Supplemental Figure 3H), one of the adverse conditions that amastigotes encounter inside human macrophages (DA SILVA et al., 2017). Thus far, based on phenotypic studies, the lack of ncRNA97 suggests its involvement in parasite duplication, triggering alterations correlated with the nutritional stress response and cell differentiation, which are essential processes for parasite survival.

One possible role for ncRNAs is the modulation of the levels of neighboring genes; these transcripts are retained at the chromatin of the locus of origin as *cis*-regulatory ncRNAs (BUMGARNER et al., 2009; ENGREITZ et al., 2016; GIL; ULITSKY, 2020). In our study, downregulation of the upstream gene (FtsX-like) was observed by RT‒qPCR in the ncRNA97^KO^ parasites (Figure 2A), and recovery to the parental levels was observed in ncRNA97^AB^ cells (Figure 2C), indicating a positive correlation between the levels of this transcript and the presence of ncRNA97 in the cells. The subcellular localization of the FtsX-like transcript was not affected by the presence or absence of ncRNA97 (Figure 2B), despite the effects on upstream gene levels. Based on the read density of ncRNA97 revealed by RNA-seq (Figure 1A), the ncRNA97 sequence could act as a 3’UTR *cis*-element and a partner of RNA binding proteins involved in the control of mRNA abundance or translation rate. However, the ectopic add-back system suggests that ncRNA97 behaves as an individual regulatory ncRNA, directly acting on its original locus, and not as a 3’UTR element of the upstream gene.

Despite its *cis*-acting role as a factor regulating FtsX-like transcript levels, RNA-seq analysis revealed that ncRNA97 also acts at the transcriptional level to modulate genes from different loci. Indeed, several transcripts were found to be up- or downregulated in ncRNA97^KO^ parasites (Figure 2D), including the core histone gene and some genes encoding amastin (Figure 2E), an amastigote-specific protein (WU et al., 2000; ROCHETTE et al., 2005). Additionally, the gene encoding the stress-induced STI1 protein was downregulated in the ncRNA97^KO^ cells, which might be linked to their lower nutritional stress resistance (Figure 4B). This protein was characterized in promastigotes from *L. major* as a heat stress response-dependent protein, in addition to complexing with Hsp70 and Hsp83 chaperones in other organisms (WEBB et al., 1997). Additionally, STI1 is present at higher levels in epimastigotes from *T. cruzi* under nutritional stress, and impaired differentiation was observed in the STI1^KO^ parasite (SCHMIDT et al., 2018). Similar stress response and metacyclogenesis phenotypes were observed for *L. braziliensis* ncRNA97^KO^ promastigotes. Consistently, the levels of these up- and downregulated genes recovered to the parental levels in ncRNA97^AB^ parasites, confirming the correlation between the levels of these genes and ncRNA97 (Figure 2F and G). Overall, the phenotypic and transcriptomic results corroborate the involvement of ncRNA97 at different stages of *Leishmania* development.

It is well established that ncRNAs might act as regulators at different levels, including through modulation of the stability or translation rate of their target mRNAs (HUANG et al., 2022). In *T. brucei,* coding and ncRNA interactions were confirmed for *Grumpy,* a ncRNA involved in parasite differentiation (GUEGAN et al., 2022). Our results reveal a correlation of ncRNA97 levels and other coding genes in the cells from distant genomic loci, suggesting that ncRNA97 may exert a *trans*-regulatory function. Thus, we may hypothesize that ncRNA97 increases the stability of its target mRNAs, particularly amastigote-specific genes, given that ncRNA97 is preferentially expressed in the intracellular stage, but the regulatory activity of ncRNA97 may involve RNA-protein interactions.

Thus, considering that such interactions might involve proteins, we conducted an *in vitro* assay to identify proteins that specifically bind to the ncRNA97 sequence. Among the 25 proteins identified with a *p* value <0.05 that also had a predicted function, we identified a putative rRNA methyltransferase protein (A4HFX5_LEIBR). The rRNA modification pathway is poorly characterized in trypanosomatids, and this methyltransferase protein is the sole protein identified to be involved in this pathway (Supplemental Figure 7B). The rRNA methyltransferase protein is a nucleus-preferential protein involved in rRNA maturation and the biogenesis of ribosomal subunits in other eukaryotes (SATTERWHITE; MANSFIELD, 2022). ncRNA97 was primarily detected in the nuclei of promastigotes in the log phase of growth (Figure 3A), which supports the hypothesis that it interacts with these macromolecules *in vivo*. rRNA biogenesis in trypanosomatids is known to be mediated by small nucleolar RNAs (snoRNAs) located mainly in the nucleus, which drive specific modifications of these rRNA molecules (CHIKNE et al., 2019). Our pulldown results indicate a putative role for ncRNA97 in mediating rRNA processing through protein interactions. Consistent with this hypothesis, an increase in noncomplexed RS was observed in ncRNA97^KO^ parasites compared to control cells (Figure 3C). Additionally, after starvation, the accumulation of ncRNA97 at a sucrose density corresponding to the pre-polysome fraction was observed (Figure 3D). In *T. brucei,* it was reported that an abundant tRNA-derived ncRNA is produced under starvation conditions and is involved in facilitating the recovery of mRNA loading to normal levels once starvation is halted (FRICKER et al., 2019). In *L. braziliensis*, despite the decrease in total ncRNA97 under nutritional stress conditions (Figure 4C), a starvation time-dependent increase in cytoplasmic ncRNA97 (∼80 times) was observed (Figure 4E), which matches its accumulation in ribosomes (Figure 3D). Thus, we may assume that ncRNA97 biogenesis is not modulated by starvation, as observed for the tRNA fragment in *T. brucei* (FRICKER et al., 2019), but that its subcellular localization is altered under this environmental condition.

Despite ncRNA97 nuclear localization, the identification of the ribosomal protein (RP) L31 and the translational initiation factor 2 (eIF2α) in the pulldown assay suggested a possible cytoplasmic localization for ncRNA97 (Supplemental Figure 7B). In *Leishmania* parasites, the level of eIF2α is directly associated with the protein synthesis rate (LAHAV et al., 2011), and eIF2α phosphorylation is essential for parasite survival under stress conditions (ABHISHEK et al., 2017), which is consistent with the poorer response of ncRNA97^KO^ parasites to nutritional stress (Figure 4B). Additionally, the interactions predicted *in vitro* suggest the relevance of ncRNA97 for ribosome functionality since RPL31 is a component of the large RS in trypanosomes and is associated with small rRNAs (srRNA3) involved in the process of ribosome assembly (LIU et al., 2016). In fact, ncRNA97 is preferentially expressed in amastigotes and seems to regulate the levels of amastigote-specific genes. However, the observed phenotypes indicate that the function of ncRNA97 does not depend on its level in the cell, since most of the alterations were observed in the promastigotes, such as a decrease in the nutritional stress response and differentiation rate (Figure 4B and D).

Ultimately, we propose both potential *cis-* and *trans-*regulatory functions for ncRNA97, coordinating the modulation of the levels of some coding genes and phenotypic alterations (Figure 5). Its putative interaction with a conserved motif in these transcripts was strongly supported by the direct effect on their levels in ncRNA97^KO^ and ncRNA97^AB^ parasites (Figure 2F and G). Together, the identified ncRNA97-interacting proteins are associated with rRNA processing and are essential for a better response to nutritional stress. Clearly, further investigation is required to fully understand and elucidate the mechanism of action of ncRNA97 and the relevance of its secondary structure, but we propose that it acts at the *cis-* and *trans-* levels of regulation. Similarly, the long ncRNA *Jpx* was identified as both a *cis-* and *trans-*regulatory element that activates *Xist,* a ncRNA that coordinates gene products *in vivo* (CARMONA et al., 2018). Finally, the results herein present the first robust molecular and functional characterization of an ncRNA in *L. braziliensis*. Other differentially expressed ncRNAs in these parasites may also play relevant roles in protozoan development, and investigating them could provide insights into ncRNAs as targets for therapeutic options in the future.

**Figure 5.**
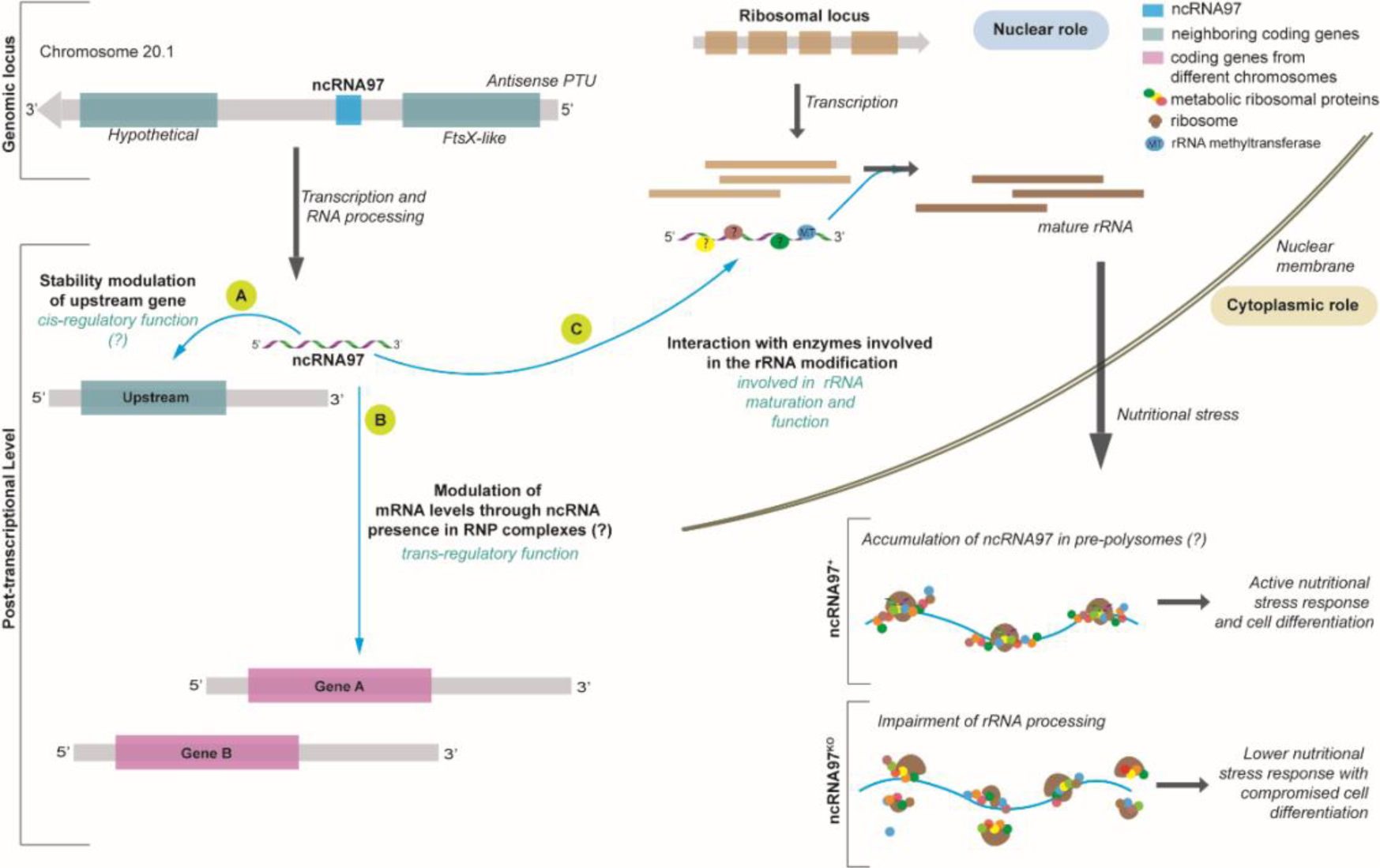
Model of the ncRNA97 levels of action in *L. braziliensis*. (A) After transcription, ncRNA97 acts as a cis-regulatory ncRNA regulating the FtsX-like transcript level. (B) ncRNA97 may also act as a trans-element regulating the levels of a group of mRNAs from different genomic loci by an unknown mechanism. Such interactions probably occur in RNP complexes that include RNA binding proteins (?). (C) Still in the nucleus, ncRNAs seem to interact with proteins involved in ribosomal (r) RNA processing, such as rRNA methyltransferase protein (MT), to generate mature rRNAs. (D) In the cytoplasm, under nutritional stress, the ncRNA97KO parasites are less resistant and exhibit a lower differentiation rate, probably due to incorrect rRNA processing. In control cells, ncRNA accumulates in the PP fraction under nutritional stress (?).

## MATERIAL AND METHODS

### Cell culture, *in vitro* differentiation, and infection assays

Promastigotes of the *Leishmania braziliensis* strain M2903 (MHOM/BR/75/M2903) expressing tdTomato fluorescent protein (LORENZON et al., 2022) were cultured in M199 medium (Sigma-Aldrich) supplemented with 10% heat-inactivated fetal bovine serum (FBS) as previously described (KAPLER; COBURN; BEVERLEY, 1990). For the metacyclogenesis assay, 10^5^ parental and knockout cells·mL-1 were incubated in 10 mL of M199 medium and grown for 120 h. Then, the metacyclic fraction was enriched by Ficoll, as described (SPÄTH; BEVERLEY, 2001). The total number of cells before purification was set as 100%, and the percentage of the metacyclic form recovered was used to estimate the differentiation rate. Purified metacyclic parasites were used to infect THP-1 human macrophages *in vitro*. THP-1 monocytes were cultivated in RPMI medium at 37°C in a 5% CO_2_ atmosphere and differentiated inside macrophages in 96-well black flat-bottom plates at 2·10^4^ cell·mL^-1^ according to an established protocol (DAIGNEAULT et al., 2010). After differentiation, the purified metacyclic forms were incubated at a ratio of 10:1 with macrophages for 4 h. Afterward, the cells were double-washed with PBS and incubated with RPMI medium for 48 h at 33°C in a 5% CO_2_ atmosphere. The nuclei were stained with 0.25% Hoechst (Merck-Millipore), and images were taken for 9 fields of view in each well under 40x magnification by the ImageXpress Micro XLS System (Molecular Devices, LLC, USA) using DAPI and Texas Red filters. The infectivity rate (%) and intracellular amastigote growth were obtained as described in a previous publication (LORENZON et al., 2022). To obtain axenic amastigotes for the oxidative stress assay, 10^5^ metacyclic promastigotes purified from the Ficoll assay were incubated in 5 mL of 100% FBS and incubated at 33°C in a 5% CO_2_ atmosphere for 72 h. Afterward, one passage of the axenic amastigotes was performed before using the cells for assays. Stage-specific markers were tested for both metacyclic and amastigote morphologies as described previously (RUY et al., 2019).

### Transfection to generate knockout and add-back parasites

The production of single guide RNA and donor DNA templates for knockout parasite generation followed the protocol from Beneke et al., 2017. After DNA recovery by ethanol precipitation, promastigotes in the log phase were transfected as previously described (LORENZON et al., 2022). Knockout parasites were selected in the presence of puromycin as the selection marker, and edited homozygotes were confirmed by conventional PCR (see Supplemental Figure2B). For add-back parasites, a plasmid was used to clone the ncRNA97 sequence (see Supplemental Figure8). Then, 10^7^ ncRNA97^KO^ parasites were transfected with 30 µg of plasmid and selected under drug pressure on solid culture media. Approximately 15 individual parasite colonies were collected after transfection, and plasmid insertion was confirmed by conventional PCR targeting the drug resistance region (streptothricin acetyltransferase – SAT).

### RNA extraction and RNA-seq data generation

Total RNA was extracted from parasites of different morphologies with a Direct-zol extraction kit (Zymo Research). RNA samples were quantified by UV‒Vis, and the quality was evaluated with an Agilent Bioanalyzer system (RNA 600 Nano kit – Agilent, Waldbronn, Germany) according to the manufacturer’s instructions. Then, RNA samples were subjected to qPCR and circularization assays. For RNA-seq data generation, total RNA from procyclic promastigotes in biological triplicates was rRNA depleted using a Ribo-Zero Plus rRNA Depletion Kit (Illumina). RNA sequencing was carried out by BGI (https://www.bgi.com/global) using a DNBSEQ Eukaryotic Strand-specific mRNA Library Preparation Kit (poly(A) tail capture and strand-specific) and the DNBseq sequencing platform (paired-end reads). After data pretreatment conducted by BGI (adapters and low-quality read removal), approximately 70 million reads per library remained. Reads were mapped to the *Leishmania braziliensis* MHOM/BR/75/M2903 genome (TriTrypDB version 56, https://tritrypdb.org/tritrypdb/app/) with Bowtie2 (version 2.3.2, (LANGMEAD; SALZBERG, 2012)) and the additional parameters *–local* and *-N 1*. Next, reads mapped to each gene locus were counted with HTSeq-count (version 0.12.4, (PUTRI et al., 2022) using the additional parameters -*-nonunique=all*, *--stranded=reverse*, *--order=pos* and *--mode=union*. Differential gene expression analysis was conducted using the R Bioconductor DESeq2 package (version 1.20.0, (LOVE; HUBER; ANDERS, 2014), and genes were considered differentially expressed when *p-value adjusted* < 0.05. An additional fold-change cutoff (FC > 1.5) was applied to prioritize genes with greater differences in expression levels between groups.

### Circular RNA assay

Total RNA extracted from log phase promastigotes was used for the determination of the 5’ and 3’ ends of the ncRNA97 transcripts following a previously published protocol (HANG et al., 2015b). Briefly, 1 µg of total RNA was treated or not treated with 1,000 U of tobacco acid pyrophosphatase (TAP) at 37°C for 1 h to generate 5’-monophosphate RNAs. Then, RNA circularization was performed using 3,000 U of T4 RNA ligase for 2 h at 37°C. Total RNA was precipitated with ethanol and recovered in 70 µL of RNAse-free water. This precipitated RNA was used to generate complementary DNA (cDNA) by reverse transcription using a gene-specific primer (cRT), followed by amplification of the region of interest by conventional PCR using specific primers for the ncRNA97 sequence (cF1/cR1 and cF2/cR2), which was directed outward of the transcript in each extremity. The amplicon was purified and cloned and inserted into a pGEM-T Easy vector. After RNA circularization, colony PCR was performed on 5 colonies for each amplicon, and the products of different sizes were sequenced. The results were analyzed with Geneious Prime Software. Details about the primer sequences, colony PCR gel, and transcript sequences are provided in Supplementary Figure 1.

### Reverse Transcription (RT) Quantitative (q)PCR

After RNA extraction, total RNA was treated with DNase Turbo (Thermo Fisher Scientific), and an RT‒qPCR assay was performed as previously described (RUY et al., 2019). The data were analyzed according to the ΔΔCt method (PFAFFL, 2001) using the geometric mean of two selected housekeeping genes (G6PD or 7SL) for normalization according to a previously described strategy (VANDESOMPELE et al., 2002). The RT‒qPCR data were analyzed using GraphPad Prism 5 (Prism). The data shown correspond to the means and standard deviations (± SDs) from 3 independent experiments. The primers used in this study were as follows: ncRNA97 Fw 5’-GTTGGTAATCGTGTGAGTGTGTG-3’, Rv 5’-GTGGTGTTTGGTAGGACGGT-3’; Amastin (LBRM2903_080014000) Fw: 5’-CCTTGCGCTCAACATCACTG-3’, Rv: 5’-AACCAGGCCACCACAAACAT-3’; Amastin (LBRM2903_100024300) Fw: 5’-CCTTCTTGGGAGTCGGCTTC-3’, Rv: 5’-CTTCTACTTCCCCTCGCTGC-3’; Stress-induced (LBRM2903_080016600) Fw: 5’-AGGGAGTGAAAGACGTGCAG-3’, Rv: 5’-TCAGGTTGCAGCATCAGGAG-3’; core histone (LBRM2903_3500053000) Fw, 5’-CCGCCGTGACTGTCTTCTT-3’; Rv, 5’-GGTGGTGTAAAGCGCATCTC-3’; and DEAD/DHA helicase (LBRM2903_3200217000) Fw 5’-CATACACGAGCGACACCGTA-3’, Rv 5’-CGTGTGGAGTCCGTGAAGAA-3’.

### Exonuclease treatment

Total RNA treated with DNase Turbo (Thermo Fisher Scientific) was incubated with the terminator 5’-phosphate-dependent exonuclease T5’ (Biosearch Technologies) to degrade 5’-monophosphate transcripts. For that, 1 µg of RNA was treated with 1 U of T5’ at 30°C for 2 h. Then, RNA was precipitated with ethanol and used for cDNA generation by reverse transcription. The presence of the ncRNA97 transcript was confirmed by conventional PCR and detected on a 1.5% agarose gel. For the positive control, genomic DNA was used. An RNA sample not subjected to reverse transcription was used as a negative control to confirm DNA degradation by DNase. The sequences of the primers used were 5’-3’ Fw_GTTGGTAATCGTGTGAGTGTGTG and Rv_GTGGTGTTTGGTAGGACGGT.

### Secondary RNA structure determination

Secondary structures of the single-stranded RNA were predicted by the RNAfold WebServer (GRUBER et al., 2008) with the minimum free energy (MFE) and partition function algorithms from the Turner model (MATHEWS et al., 2004). All sequenced transcripts for ncRNA97 found in the circularization assay were subjected to secondary structure prediction.

### Pulldown assay – *in vitro* RNA‒protein interaction

The sequences corresponding to the predicted ncRNA97 sequence were amplified from the genomic DNA using the primers Fw (5’-CGTTGTTGGTAATCGTGTGAGT-3’) and Rv (5’-AGTGGTGTTTGGTAGGACG-3’). Then, it was cloned and inserted into a pUC-56 plasmid between the T7 promoter and 4xS1m aptamer sequences, adapted from Leppek’s strategy (26). RNA was transcribed *in vitro* (MEGAscript T7 transcription kit – Thermo Fisher AM1334), and 10 µg of purified RNA was immobilized on streptavidin magnetic beads (NEB) for 8 h at 4°C under orbital rotation in binding buffer (24 mM KCl, 10 mM Tris-base pH 7.8, and 10 mM MgCl_2_). All experiments were performed in an RNAse-free environment with diethylpyrocarbonate (DEPC) pretreatment and the addition of RNAse inhibitors (200 U). An empty sequence between the T7 promoter and 4xS1m aptamer was used as an RNA control to identify nonspecific partners. A total of 10^8^ parasites were lysed in 1 mL of SA-RNP-Lyse buffer (20 mM Tris-HCl pH 7.5, 150 mM NaCl, 1.5 mM MgCl_2_, 2 mM DTT, 2 mM RNase inhibitor, 1 protease inhibitor cocktail tablet (Merck), 1% Triton X-100) on ice with physical pressure using a 19 G needle. Biotinylated proteins were removed from the extract by incubating the lysed extract for 8 h at 4°C with streptavidin beads. Subsequently, the supernatant was incubated with the bead-immobilized wild-type ncRNA97 sequence for 8 h at 4°C. Afterward, the beads were washed three times with wash buffer (20 mM Tris-HCl pH 7.5, 300 mM NaCl, 5 mM MgCl2, 2 mM DTT, 2 mM RNase inhibitor, and 1 protease inhibitor cocktail tablet), resuspended in 35 µL of Laemmli buffer (LAEMMLI, 1970) and boiled for 10 min before being applied to a 12% polyacrylamide gel. Electrophoresis was run at 110 V until the samples reached the separation gel.

### Proteomic analysis, mass spectrometry, database search, and protein identification criteria

Gel bands from the samples were sent for protein identification by mass spectrometry analysis (Proteomics Platform of the CHU de Quebéc Research Centre, Quebec, Canada). Three biological replicates were evaluated for the ncRNA97 sequence and the control. The results were obtained and analyzed using Scaffold Protein software. The list of identified proteins was filtered using a protein threshold of 99%, a peptide threshold of 95%, and a minimum of 1 peptide identified in each sample. Proteins interacting with the control RNA sequence were excluded, and only proteins that specifically interacted with the ncRNA97 sequence in triplicate samples were included in the results. The details of the mass spectrometry and database search protocols and criteria for protein identification have been described elsewhere (LORENZON et al., 2022).

### Phenotypic assays

To evaluate the nutritional stress response of promastigotes in the log phase, parasites at 10^6^ cells·mL^-1^ were incubated for 2 or 4 h in PBS for total starvation. Subsequently, the cells were pelleted and resuspended in 100 µL fresh M199 medium and seeded in a 96-well plate for 20 h at 27°C for recovery. Then, 10 µL of 1 mg·mL^-1^ MTT was added to each well, and the plate was incubated for another 4 h. Afterward, the plate was centrifuged at 2,000 × g, and the pellet was resuspended in 100 µL of DMSO, after which the absorbance was measured at 570 nm. Non-starved cells were used as a negative control and were considered to have 100% cell viability. For oxidative stress response assays, 10^6^ cells·mL^-1^ axenic amastigotes were incubated for 24 h in FBS containing 300 µM H_2_O_2_at 33°C in a 5% CO_2_ atmosphere. After treatment, the cells were pelleted, resuspended in 100 µL fresh FBS, and then seeded in a 96-well plate for the same viability assay as described for nutritional stress.

### Polysome profile analysis

Promastigote polysomes were analyzed under normal and nutritional stress conditions by purification and fractionation on a sucrose gradient (Holetz et al., 2007). Briefly, 10^9^ cells were starved for 2 h in 100% PBS, and then 100 µg·mL^-1^ cycloheximide was added for 10 min. On ice, the cells were washed once with TKM buffer supplemented with 100 µg·mL^-1^ cycloheximide, 10 µg·mL^-1^ heparin, 10 µg·mL^-1^ E-64 and protease inhibitor cocktail for polysome blocking (Roche). For polysome dissociation, cells were treated with 2 mM puromycin. The cells were pelleted, 100 µL of buffer lysis buffer (TKM supplemented with 10% surfactant NP-40 and 2 M sucrose) was added, and the cells were lysed by up-and-down agitation followed by centrifugation at 18,000 × g at 4°C for 10 min. The supernatant was layered on top of a linear 10-50% sucrose density gradient prepared in the same buffer. The system was centrifuged at 39,000 rpm at 4°C for 2 h in a Beckman SW41 rotor. After centrifugation, the gradient fractions were collected with an ISCO gradient fractionation system, and 500 µL of each collected sample was used for RNA extraction and qPCR.

### Statistical analysis

The experiments were performed in biological triplicates, and statistical analyses (ANOVA with Dunnett’s test) were performed using GraphPad Prism 8. *p*<0.05 was considered to indicate statistical significance. Statistically significant differences are indicated by asterisks, where \*\**p*≤ 0.002, \*\*\**p*≤ 0.0002 and \*\*\*\**p*≤ 0.0001.

## ACKNOWLEDGMENTS

We would like to thank Susanne Kramer from the University of Würzburg, Germany, for the critical discussions that improved the work and for opening her laboratory and resources for training for J.C.Q.J.; Viviane Ambrósio for the technical support in the laboratory; Gustavo D. Campagnaro for sharing tools; Pegine Walrad from the University of York for materials exchanges; and the staff of the Proteomics Platform of the CHU de Québec Research Centre, Québec, Canada. We also acknowledge TriTrypDB for providing indispensable resources and tools.

